# Genetics of Fertility Restoration in A_1_ Cytoplasmic Genetic Male Sterility (CGMS) Systems in Sorghum (*Sorghum bicolor* L. Moench)

**DOI:** 10.1101/2022.03.23.485454

**Authors:** Gopal W Narkhede, Shivaji P Mehtre, Krishnananda P Ingle, Kirandeep Kaur Romana, KS Vinutha, Santosh P Deshpande

## Abstract

The cause of Cytoplasmic Genetic Male Sterility (CGMS) is specific nuclear and mitochondrial interactions. Almost all commercial sorghum hybrids were developed using the A1 cytoplasmic genetic male sterility system. Understanding the inheritance of fertility restoration in sorghum for A1 cytoplasm, for example, can improve the selection efficiency of restorer lines for increased seed production. In a cross of male sterile line 296A with A1 cytoplasm and restorer lines comprised of a set of Recombinant Inbred Lines (RILs), the inheritance pattern of fertility restoration of sorghum was studied. The F_1_ hybrid was completely fertile, revealing the dominant nature of fertility restoration, which is controlled by one or two major genes with modifiers. In this study, the genetics of fertility restoration of the A1 cytoplasmic nuclear male sterility system (CGMS) in sorghum were investigated in segregating F_2_ and BC_1_ populations of A1 cytoplasm crosses. Fertility restoration was governed by a monogenic inheritance (3F:1S) mechanism represented by a single dominant gene responsible for fertility restoration in all of the crosses studied.

## 1. Introduction

Sorghum originated in Africa and evolved into an important cereal crop. It now feeds over 500 million people in 98 countries, with 42 million hectares of cultivated land and 62 million tonnes of yields per year. The global sorghum crop area is 39.93 million hectares, with a total production of 59.35 million tonnes and a productivity of 6.7 metric tonnes per hectare (FAOSTAT). The interaction of sterility-inducing factors in the cytoplasm with genetic factors in the nucleus causes cytoplasmic-genetic male sterility (CGMS). This system focuses on three lines of breeding: A-, B-, and R-lines. Because most CMS lines have irregular anther or pollen formation, the A-line is a male sterile. When the B- and A-lines were crossed, they produced genetically male sterile offspring. The mean difference between A-line and B-line deals with flowering. The third line is called the restorer line (R-line). The restorer line’s purpose is to serve as a pollinator variety for pollinating the CMS line in order to produce F1 hybrids (Yuan et al. 2003). It is essential for the restorer line to have strong restoring ability. To ensure successful pollen transfer from R-line to A-line, the restorer line should have venerable combining ability and agronomic characteristics, as well as a well developed flowering system. Stephens and Holland (1954) discovered A1 cytoplasmic genic male sterility (CGMS) in sorghum and exploited it for hybrid production worldwide. The genetic makeup of the cytoplasm and nuclei determines the inheritance of male sterility/fertility. In some cases, single genes control male fertility restoration, but it is polygenic when the same nuclear genotype interacts with different cytoplasm. Intra and inter-allelic interactions and complementation influence fertility restoration. The thorough understanding of the genetics of fertility restoration is useful in planning a sound breeding strategy for the development of superior restorers in a hybrid breeding program. It may also help in the efficient transfer of restorer genes into other agronomical desirable genotypes. Cytoplasmic male sterility which causes the production of non-functional pollen and inherited maternally is important in commercial hybrid seed production and breeding program. In last few decades, productivity of field, vegetables and fruit crops was increased due to this hybrid breeding technique. Efficiency of exploitation of heterosis at commercial level increased ue to availability of more number of cytoplasmic genetic male sterile sources for fertility restoration. Dominant fertility restoring nuclear genes are transmitted from the male parent, which allow seed set on the hybrid plants. However, the expression of fertility restoration may vary from 0 % to 100 % fertility restoration of CGMS-based hybrids is an integral part of breeding hybrids and the development of new hybrid parents with desirable agronomic and market preferred traits on regular intervals is essential for the sustainability of hybrid technology programs. The presence of greater genetic diversity among fertility restorers enhances the probability of breeding widely adapted high yielding hybrids (Dandin et al. 2014; Praveen et al. 2015). The information about the number of genes controlling fertility restoration in the nucleus suppresses the male-sterile phenotype and allows commercial exploitation of CMS system for the production of hybrid seeds. Therefore, the present investigation was undertaken to assess the genetics of fertility restoration system in sorghum using F_2_ and BC_1_F_1_ generations in three sorghum hybrids carrying A1 cytoplasm.

## 2. Materials and Methods

Cytoplasmic genetic male sterile line (296A) crossed with 238 RILs (Recombinant Inbred Lines) individuals (296B x IS 18551) to produced F_1_ hybrids, evaluated in rainy and post-rainy season 2016 for seed setting percentage and pollen fertility/viability score. From these multi-season evaluation program, results revealed that three F_1_’s *viz;* 296A x ICSL 43119, 296A x ICSL 43123 & 296A x ICSL 43126 were selected to study genetics of fertility restoration on the basis of high seed set percent (> 90 %) and high pollen fertility/viability score (8-9 score). All these selected F_1_ plants were selfed to produce F_2_ seeds and simultaneously crossed all the F_1_ plants to their male sterile line (296A) to produced BC_1_ seeds. The F_2_’s and BC_1_s along with their parents were raised at International Crops Research Institute for Semi-Arid Tropics, Patancheru, India during late post-rainy season 2017-18. Prior to flowering, heads from each line were covered with paper bags to exclude foreign pollen. At 40 days after the crop flowered, the bags were removed and the percent seed set on each head was visually rated. On the basis of seed setting percentage the F_2_ and BC_1_ classified into fertile, partial fertile and sterile (Prasad and Biradar, 2018). The F_2_ and BC_1_ populations were tested for segregation ratios to determine the number of genes involved in the fertility restoration of cytoplasmic genetic male sterility (CGMS) systems. The goodness of fit to the expected ratios in F_2_ and BC_1_F1 generations were tested using chi-square test (Meshram and Patil, 2018; Praveen et al., 2015, Arunkumar et al., 2004; Dandin et al., 2014).

## 3. Statistical Analysis

All the data analysis was carried out in R software version 3.6.3 (R Core Team, 2019). Chi square (χ2) method with Yates correction factor (**Steel and Torrie, 1980**) was applied on the observed data to test the goodness of fit of different genetic ratios. The calculated χ2 values were compared with the tabulated χ2 values with (n-1) degrees of freedom at 5 % and 1 % probability level. The null hypothesis was rejected if the calculated χ2 value exceeded the corresponding tabulated χ2 value. Exact probability value at (n-1) degrees of freedom for the best fit hypothetical ratio was calculated in the Excel spreadsheet using the statistical function ‘CHIDIST’.

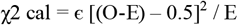

Where,

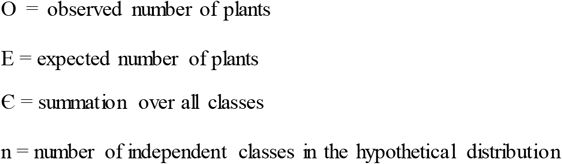

## 4. Results

The inheritance of A_1_ CMS system was investigated on F_2_ and BC_1_ in a total of 3 (A x R) crosses. All the F_2_s and BC_1_F_1_s were evaluated at ICRISAT-Patancheru for seed setting data. In crosses where seed setting data was recorded, genetic ratios were worked out by distribution of plants in their respective groups *viz;* seed setting percentage between 0-10 % classified as sterile, seed setting percentage between 11-60 % were grouped as partial sterile while seed setting percentage between 61-100 % were considered as fertile (Prasad and Biradar, 2018). The results of inheritance of A_1_ CMS system with respect to each of the fertility restorer parents have been presented below.

In the present study, In cross 296A x ICSL 43119, 610 F_2_ plants segregated in 458 fertile (F) and 152 sterile (S) plants and had a good fit (χ2 = 0.0; P = 0.963) to the ratio of 3F:1S which revealed single dominant gene responsible for fertility restoration. 590 BC_1_F_1_ plants of the same distributed in 299 fertile (F) and 291 sterile (S) and show segregation of 1F:1S (χ2 = 0.11; P = 0.742). In cross 296A x ICSL 43123, 595 F_2_ plants segregated in 430 fertile (F) and 165 sterile (S) and fit to the hypothesized 3F:1S ratio (χ2 = 2.37, P = 0.124). In the same way, 585 BC_1_F_1_ plants divided in 301 fertile (F) and 284 sterile (S) show segregation according to ratio of 1F:1S as indicated by χ2 value of 0.49 (P = 0.482). In cross 296A x ICSL 43126, 495 F_2_ plants segregated in 375 fertile (F) and 120 sterile (S) had a good fit (χ2 = 0.15; P = 0.697) to 3F:1S while, 400 BC_1_F_1_ plants segregated in 208 fertile (F) and 192 sterile (S) was in good agreement with the expected ratio of 1F:1S as evident from χ2 value of 0.64 (P = 0.424) (Table 1).

**Table 1:** Segregation ratios for fertile and sterile plants in F_2_ and backcross population derived from crosses based on A_1_ cytoplasm based A-lines in sorghum. (Monogenic Interaction)

| SN | Cross | Generations | No of Plants | | | Expected ratio (F:S) | $\chi^2$ value | Probability value (P) |
| --- | --- | --- | --- | --- | --- | --- | --- | --- |
|  |  |  | Total | Fertile | Sterile |  |  |  |
| **1** | **296A x ICSL 43119** | **P<sub>1</sub>** | 10 | 0 | 10 |  |  |  |
|  |  | **P<sub>2</sub>** | 9 | 9 | 0 |  |  |  |
|  |  | **F<sub>2</sub>** | 610 | 458 | 152 | 3:1 | 0.00 | 0.963 ns |
|  |  | **BC<sub>1</sub>F<sub>1</sub>** | 590 | 299 | 291 | 1:1 | 0.11 | 0.742 ns |
| **2** | **296A x ICSL 43123** | **P<sub>1</sub>** | 10 | 0 | 10 |  |  |  |
|  |  | **P<sub>2</sub>** | 9 | 9 | 0 |  |  |  |
|  |  | **F<sub>2</sub>** | 595 | 430 | 165 | 3:1 | 2.37 | 0.124 ns |
|  |  | **BC<sub>1</sub>F<sub>1</sub>** | 585 | 301 | 284 | 1:1 | 0.49 | 0.482 ns |
| **3** | **296A x ICSL 43126** | **P<sub>1</sub>** | 10 | 10 | 0 |  |  |  |
|  |  | **P<sub>2</sub>** | 11 | 0 | 11 |  |  |  |
|  |  | **F<sub>2</sub>** | 495 | 375 | 120 | 3:1 | 0.15 | 0.697 ns |
|  |  | **BC<sub>1</sub>F<sub>1</sub>** | 400 | 208 | 192 | 1:1 | 0.64 | 0.424 ns |

## 5. Discussion

Mechanisms by which male fertility restoration occurs are probably as diverse as mechanisms by which mitochondrial mutations cause CMS. Although restorer alleles are known to affect all the well-characterized CMS-associated genes, mechanism of action has not been determined definitively for any restorer allele. The knowledge of fertility restoration genetics is utmost important for the transfer of restorer genes from one genotype to another (Sawargaonkar et al. 2012). In certain cases, the environment also plays an important role in the expression of pollen fertility. The presence of homozygous recessive alleles at one locus produces partial fertility, whereas the presence of fertility restoring alleles at the other locus produces male sterility (Saroj et al., 2015).

In the present study, The A_1_ cytoplasm based Recombinant Inbred Line (RIL) hybrids were evaluated multi season for seed setting percentage and pollen fertility score. Seed setting percentage and pollen fertility score recorded in rainy season on Recombinant Inbred Line (RIL) population along with pollen fertility score recorded in post rainy season had a good fit to the monogenic expected ratio of 3:1 (Campa et al. 2014). 3 F_2_s and 3 BC_1_F_1_s developed from single crosses *viz;* 296A x ICSL 43119, 296A x ICSL 43123 and 296A x ICSL 43126 evaluated to study genetics of fertility restoration. Monogenic ratio (3F:1S) showed non-significant χ2 values concluding that observed and expected values are with very negligible and/or no differences (Ali et al., 2011; Reddy et al., 2010; Rongbai et al., 2005; Sawargaonkar et al., 2012; Sreedhar et al., 2011; Murty and Gangadhar, 1990). The results are discussed below.

In six segregating populations (3 F_2_s and 3 BC_1_F1s) developed from three crosses, a monogenic F_2_ ratio of 3F:1S and the corresponding BC_1_ ratio of 1F:1S was observed that results from single dominant gene involved in fertility restoration. Consider ‘A’ gene involved in fertility restoration of the A_1_ CMS system. The monogenic F_2_ ratio of 3F:1S and the corresponding BC_1_F_1_ ratio of 1F:1S is possible when the genotype of female parent is ‘aa’ and of restorer parents is ‘AA’ indicating that single dominant gene involved in fertility restoration. The F_1_ of these parents will be fertile and heterozygous for single locus (Aa). A plant in the F_2_ will be fertile if it possesses dominant allele of the basic gene/s. All other plants will be sterile. In backcross, the F_1_ plant (Aa) crossed with female parent (aa). From this, half plants containing dominant allele of the basic gene will be fertile and others will be sterile. Monogenic modes of inheritance have been reported in sorghum for the A_1_ *(milo)* CMS system (Maunder and Pickett, 1959; Murty and Gangadhar, 1990; Arunkumar et al., 2004; Praveen et al., 2015), for A_2_ (Praveen et al., 2015), for maldandi cytoplasm (Dandin et al., 2014) and for 9E and A_4_ CMS systems (Elkonin et al., 1998); in pearl millet for A_4_ CMS system (Gupta et al. 2012) and in A_5_ (Gupta et al. 2018) in rice for CMS-BT (Komori et al., 2003), CMS-HL (Huang et al., 2000), in sunflower CMS-PET1 (Chandra et al., 2010; Sujatha et al., 2011) and for CMS-ANL2, CMS-PEF1 and CMS-PET2 (Chandra et al., 2010; Sujatha et al., 2011), in rice (Ahmadikhah et al., 2007; Sreedhar et al., 2011), in Brassica (Ahmad et al., 2013; Bisht et al., 2015), in pigeon pea (Sawargaonkar et al., 2012).

## 6. Conclusion

It is necessary to keep upgrading the hybrid technology so that high yielding hybrids can be developed. The knowledge on the genetics of fertility restoration helps in designing a plan for breeding elite hybrid parents. The genetics of fertility restoration of the A1 cytoplasmic male sterility system was governed by a single dominant gene in this study. Crosses among diverse restorer lines are required for CMS based hybrid breeding so that genotypes with high seed setting percentage can be selected.

